# Metagenomics workflow for hybrid assembly, differential coverage binning, transcriptomics and pathway analysis (MUFFIN)

**DOI:** 10.1101/2020.02.08.939843

**Authors:** Renaud Van Damme, Martin Hölzer, Adrian Viehweger, Bettina Müller, Erik Bongcam-Rudloff, Christian Brandt

## Abstract

Metagenomics has redefined many areas of microbiology. However, metagenome-assembled genomes (MAGs) are often fragmented, primarily when sequencing was performed with short reads. Recent long-read sequencing technologies promise to improve genome reconstruction. However, the integration of two different sequencing modalities makes downstream analyses complex. We, therefore, developed MUFFIN, a complete metagenomic workflow that uses short and long reads to produce high-quality bins and their annotations. The workflow is written by using Nextflow, a workflow orchestration software, to achieve high reproducibility and fast and straightforward use. This workflow also produces the taxonomic classification and KEGG pathways of the bins and can be further used by providing RNA-Seq data (optionally) for quantification and annotation. We tested the workflow using twenty biogas reactor samples and assessed the capacity of MUFFIN to process and output relevant files needed to analyze the microbial community and their function. MUFFIN produces functional pathway predictions and if provided *de novo* transcript annotations across the metagenomic sample and for each bin.

**Author Summary:** RVD did the development and design of MUFFIN and wrote the first draft; BM and EBR did the critical reading and correction of the manuscript; MH did the critical reading of the manuscript and the general adjustments for the metagenomic workflow; AV did the critical reading of the manuscript and adjustments for the taxonomic classifications. CB supervised the project, did the workflow design, helped with the implementation, and revised the manuscript.

## Introduction

Metagenomics is widely used to analyze the composition, structure, and dynamics of microbial communities, as it provides deep insights into uncultivatable organisms and their relationship to each other ^1–5^. In this context, whole metagenome sequencing is mainly performed using short-read sequencing technologies, predominantly provided by Illumina. Not surprisingly, the vast majority of tools and workflows for the analysis of metagenomic samples are designed around short reads. However, long-read sequencing technologies such as provided by PacBio or Oxford Nanopore Technologies (ONT) retrieve genomes from metagenomic datasets with higher completeness and less contamination ^6^. The long-read information bridges gaps in a short-read-only assembly that often occur due to intra- and interspecies repeats ^6^. Complete viral genomes can be already identified from environmental samples without any assembly step via nanopore-based sequencing ^7^. Combined with a reduction in cost per gigabase ^8^ and an increase in data output, the technologies for sequencing long reads quickly became suitable for metagenomic analysis ^9–12^. In particular, with the MinION, ONT offers mobile and cost-effective sequencing device for long reads that paves the way for the real-time analysis of metagenomic samples. Currently, the combination of both worlds (long reads and high-precision short reads) allows the reconstruction of more complete and more accurate metagenome-assembled genomes (MAGs) ^6^.

One of the main challenges and bottlenecks of current metagenome sequencing studies is the orchestration of various computational tools into stable and reproducible workflows to analyze the data. A recent study from 2019 involving 24,490 bioinformatics software resources showed that 26 % of all these resources are not currently online accessible ^13^. Among 99 randomly selected tools, 49 % were deemed ‘difficult to install,’ and 28 % ultimately failed the installation procedure. For a large-scale metagenomics study, various tools are needed to analyze the data comprehensively. Thus, already during the installation procedure, various issues arise related to missing system libraries, conflicting dependencies and environments or operating system incompatibilities. Even more complicating, metagenomic workflows are computing intense and need to be compatible with high-performance compute clusters (HPCs), and thus different workload managers such as SLURM or LSF. We combined the workflow manager Nextflow^14^ with virtualization software (so-called ‘containers’) to generate reproducible results in various working environments and allow full parallelization of the workload on a higher degree.

Several workflows for metagenomic analyses have been published, including MetaWRAP(v1.2.1)^15^, Anvi’o^16^, SAMSA2^17^, Humann^18^, or MG-Rast^19^. Unlike those, MUFFIN allows for a hybrid metagenomic approach combining the strengths of short and long reads. It ensures reproducibility through the use of a workflow manager and reliance on either install recipes (Conda ^20^) or containers (Docker^21^).

## Design and implementation

MUFFIN integrates state-of-the-art bioinformatic tools via Conda recipes or Docker containers for the processing of metagenomic sequences in a Nextflow workflow environment (Figure 1). MUFFIN executes three steps subsequently or separately if intermediate results, such as MAGs, are available. As a result, a more flexible workflow execution is possible. The three steps represent common metagenomic analysis tasks and are summarized in Figure 1:

1. Assemble: Hybrid assembly and binning
2. Classify: Bin quality control and taxonomic assessment
3. Annotate: Bin annotation and KEGG pathway summary

**Figure 1:**
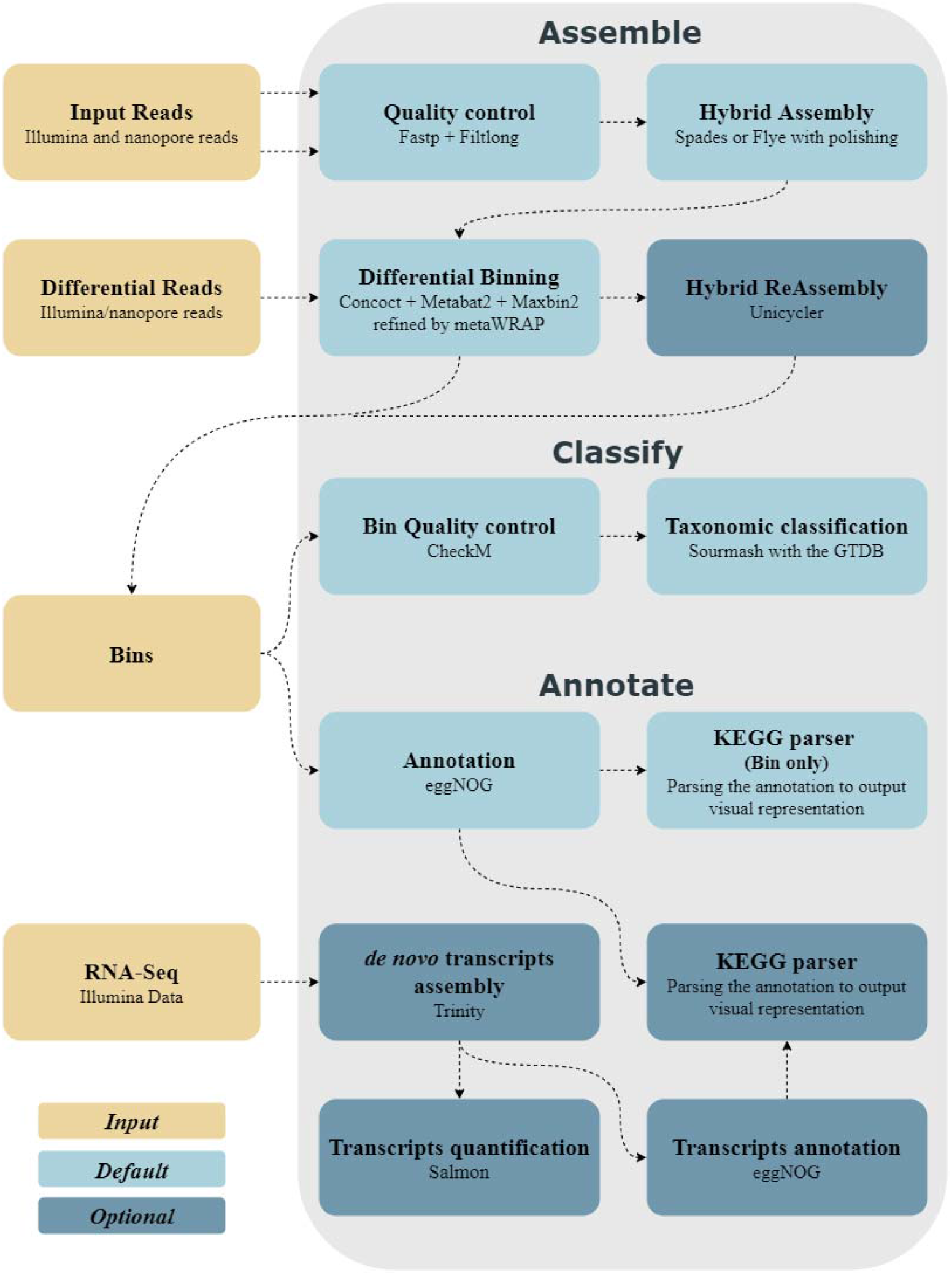
Simplified overview of the MUFFIN workflow. All three steps (Assemble, Classify, Annotate) from top to bottom are shown. The RNA-Seq data for Step 3 (Annotate) is optional.

The workflow takes paired-end Illumina reads (short reads) and nanopore-based reads (long reads) as input for the assembly and binning and allows for additional user-provided read sets for differential coverage binning. Differential coverage binning facilitates genome bins with higher completeness than other currently used methods ^22^. Step 2 will be executed automatically after the assembly and binning procedure or can be executed independently by providing MUFFIN a directory containing MAGs in FASTA format. In step 3, paired-end RNA-Seq data can be optionally supplemented to improve the annotation of bins.

On completion, MUFFIN provides various outputs such as the MAGs, KEGG pathways, and bin quality/annotations. Additionally, all mandatory databases are automatically downloaded and stored in the working directory or can be alternatively provided via an input flag.

### Step 1 - Assemble: Hybrid assembly and binning

The first step (**Assembly and binning**), uses metagenomic nanopore-based long reads and Illumina paired-end short reads to obtain high-quality and highly complete bins. The short-read quality control is operated using fastp (v0.20.0) ^23^. Optionally, Filtlong (v0.2.0) ^24^ can be used to discard long reads below a length of 1000 bp ^24^. The hybrid assembly can be performed according to two principles, which differ substantially in the read set to begin with. The default approach starts from a short-read assembly where contigs are bridged via the long reads using metaSPAdes (v3.13.1) ^25–27^. Alternatively, MUFFIN can be executed starting from a long-read-only assembly using metaFlye (v2.6) ^28,29^ followed by polishing the assembly with the long reads using Racon (v1.4.7) ^30^ and medaka (v0.11.0) ^31^ and finalizing the error correction by incorporating the short reads using multiple rounds of Pilon (v1.23) ^32^.

Binning is the most crucial step during metagenomic analysis. Therefore, MUFFIN combines three different binning software tools, respectively CONCOCT (v1.0.0) ^33^, MaxBin2 (v2.2.4) ^34^, and MetaBAT2 (v2.14) ^35^ and refine these bins via MetaWRAP (v1.2.1)^15^. The user can provide additional read data sets (short or long reads) to perform automatically differential coverage binning to assign contigs to their bins better.

Moreover, an additional reassembly of bins has shown the capacity to increase the completeness and N50 while decreasing the contamination of the bins^15^. Therefore, MUFFIN allows for an optional reassembly to improve the continuity of the MAGs further. This re-assembly is performed by retrieving the reads belonging to one bin and doing an assembly with Unicycler (v0.4.8) ^36^.

To support a transparent and reproducible metagenomics workflow, all reads that cannot be mapped back to the existing high-quality bins (after the refinement) are available as an output for further analysis. These reads could be further analyzed by other tools or, e.g., used as a new input to run MUFFIN while providing other read sets for the differential coverage binning to extract additional high-quality bins.

### Step 2 - Classify: Bin quality control and taxonomic assessment

In the second step (**Bin quality control and taxonomic assessment**), the quality of the bins is evaluated with CheckM (v1.0.18) ^37^ followed by assigning a taxonomic classification to the bins using sourmash (v2.0.0a10) ^38^ and the Genome Taxonomy Database (GTDB release r89) ^39^. The GTDB was chosen as it contains many unculturable bacteria and archaea – this allows for monophyletic species assignments, which other databases do not assure ^40,41^. GTDB substantially improved overall downstream results ^40^. The user can also analyze other bin sets in this step regardless of their origin by providing a directory with multiple FASTA files (bins).

### Step 3 - Annotate: Bin annotation and KEGG pathway summary

The last step of MUFFIN (**Bin annotation and output summary**) comprises the annotation of the bins using eggNOG-mapper (v2.0.1) ^42^ and the eggNOG database (v5) ^43^. If RNA-Seq data of the metagenome sample is provided (Illumina, paired-end), quality control using fastp (v0.20.0) ^23^ and a *de novo* transcript assembly using Trinity (v2.8.5) ^44^ followed by a quasi-mapping transcript quantification using Salmon (v0.15.0) ^45^ are performed. Lastly, the transcripts are annotated using eggNOG-mapper (v2.0.1) ^42^ again, followed by a parser to output the activity of the pathway graphically in relation to the sample level. The expression of low and high abundant genes present in the bins is shown. If only bin sets are provided without any RNA-Seq data, the pathways of all the bins are created based on gene presence alone. The KEGG pathway results are summarized in detail as interactive HTML files (example snippet: Figure 2).

**Figure 2:**
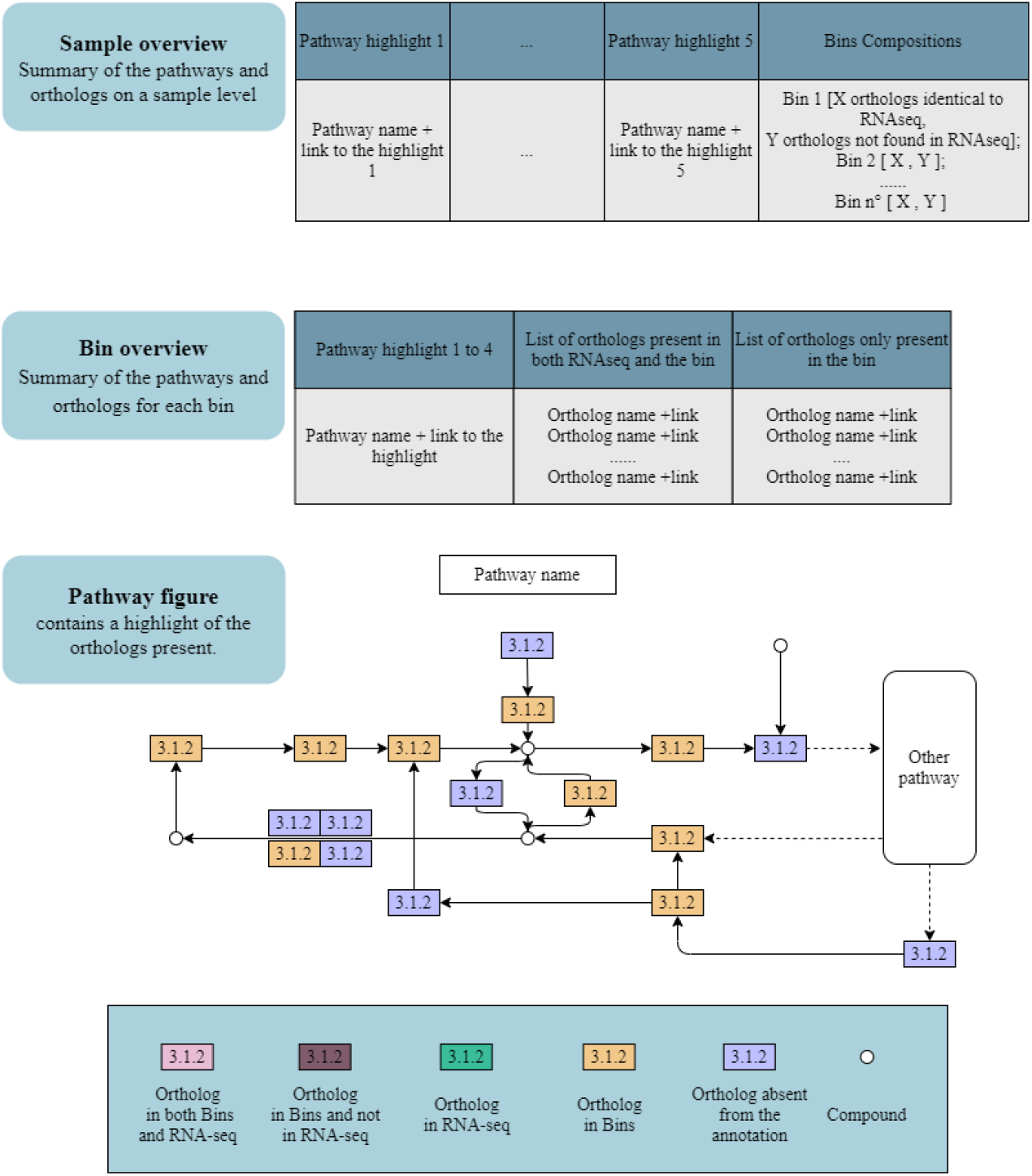
Example snippets of the sub-workflow results of step 3 (Annotate).

Like step 2, this step can be directly performed with a bin set created via another workflow.

### Running MUFFIN and version control

MUFFIN requires only two dependencies, which allows an easy and user-friendly workflow execution. One of them is the workflow management system Nextflow ^14^ and the other can be either Conda ^20^ as a package manager or Docker ^21^ to use containerized tools. A detailed Installation process is available on https://github.com/RVanDamme/MUFFIN. Each MUFFIN release specifies the Nextflow version it was tested on, to avoid any version conflicts between MUFFIN and Nextflow at any time. A Nextflow-specific version can always be directly downloaded as an executable file from https://github.com/nextflow-io/nextflow/releases, which can then be paired with a compatible MUFFIN version via the -r flag.

## Results

We chose Nextflow for the development of our metagenomic workflow because of its direct cloud computing support (Amazon AWS, Google Life Science, Kubernetes), various ready-to-use batch schedulers (SGE, SLURM, LSF), state-of-the-art container support (Docker, Singularity) and accessibility of a widely used software package manager (Conda). Moreover, Nextflow ^14^ provides a practical and straightforward intermediary file handling with process-specific work directories and the possibility to resume failed executions where the work ceased. Additionally, the workflow code itself is separated from the ‘profile’ code (which contains Docker, Conda, or cluster related code), which allows for a convenient and fast workflow adaptation to different computing clusters without touching or changing the actual workflow code.

The entire MUFFIN workflow was executed on 20 samples from the Bioproject PRJEB34573 (available at ENA or NCBI) using the Cloud Life Sciences API (google cloud) with docker containers. This metagenomic bioreactor study provides paired-end Illumina and nanopore-based data for each sample ^41^. We used five different Illumina read sets of the same project for differential coverage binning, and the workflow runtime was less than two days for all samples. MUFFIN was able to retrieve 1122 MAGs with genome completeness of at least 70 % and contamination of less than 10 % (Figure 3). In total, MUFFIN retrieved 654 MAGs with genome completeness of over 90 %, of which 456 have less than 2% contamination out of the 20 datasets. For comparison, a recent study was using 134 publicly available datasets from different biogas reactors and retrieved 1,635 metagenome-assembled genomes with genome completeness of over 50% ^46^.

**Figure 3:**
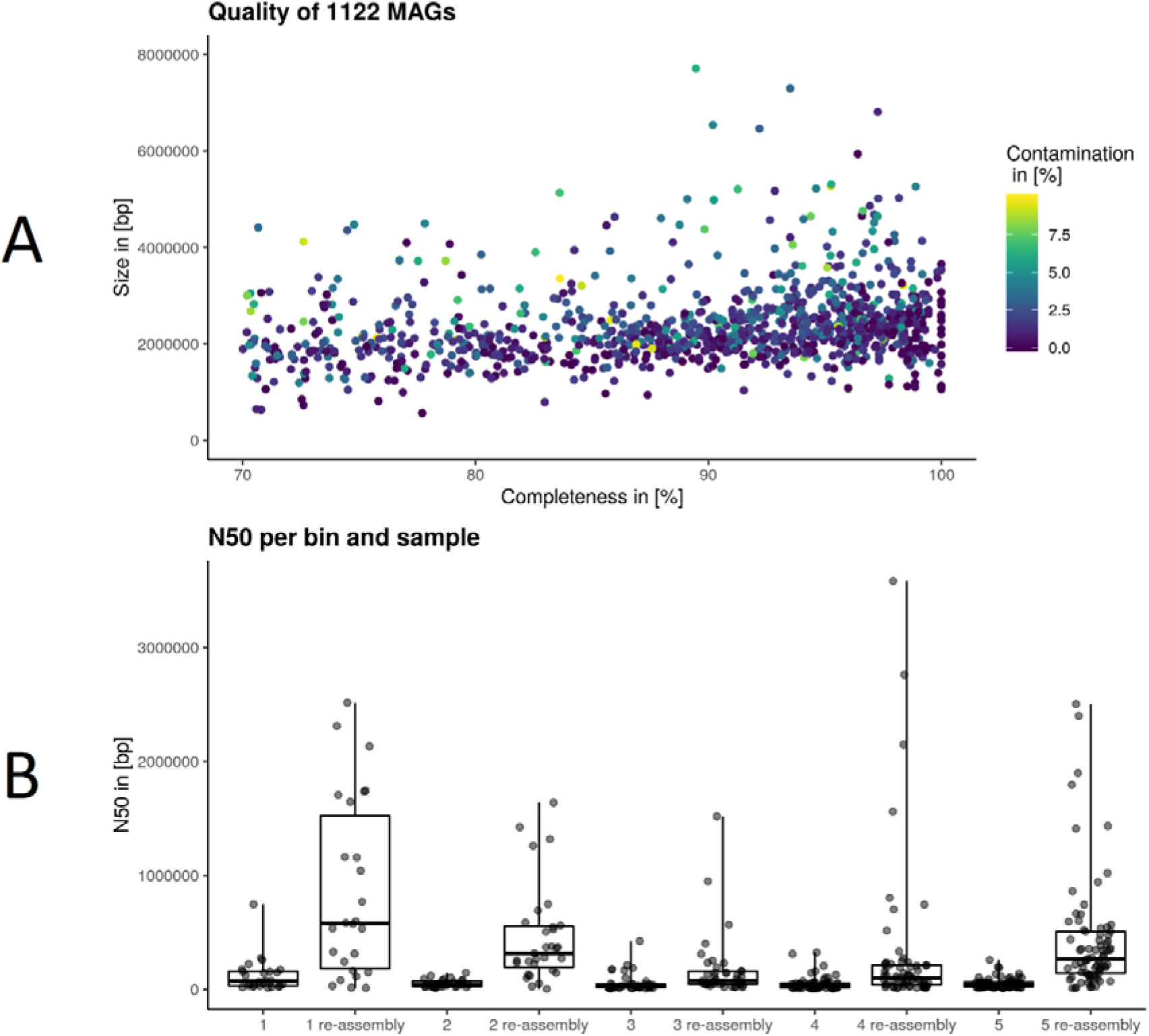
A: Quality overview of 1122 meta-assembled genomes (MAGs) by plotting size to completeness and coloring based on contamination level. B: N50 comparison between each bin of five selected samples from the Bioproject PRJEB34573 before and after individual bin reassembly.

Exemplarily, we investigated the impact of additional re-assembly of each bin for five samples (Figure 3). The N50 was increased by an average of 6-7 fold across all samples. Twenty-six bins of the five samples had an N50 ranging between 1 to 3 Mbases. Some bins benefit more of this step as the re-assembly performance depends on the number of reads available for each bin.

## Discussion

The analysis of metagenomic sequencing data evolved as an emerging and promising research field to retrieve, characterize, and analyze organisms that are difficult to cultivate. There are numerous tools available for individual metagenomics analysis tasks, but they are mainly developed independently and are often difficult to install and run. The MUFFIN workflow gathers the different steps of a metagenomics analysis in an easy-to-install, highly reproducible, and scalable workflow using Nextflow which makes them easily accessible to researchers.

MUFFIN utilizes the advantages of two sequencing technologies, whereas short reads can provide a better representation of low abundant species due to their higher coverage. This aspect is further utilized via the final re-Assembly step after binning, which is an optional step due to the additional computational burden which solely aims to improve genome continuity.

Another critical aspect is the full support of differential binning, for both long and short reads, via a single input option. The additional coverage information from other read sets of similar habitats allows for the generation of more concise bins with higher completeness and less contamination because more coverage information is available for each binning tool to decide which bin each contig belongs.

With supplied RNA-Seq data, MUFFIN is capable of enhancing the pathway results present in the metagenomic sample by incorporating this data as well as the general expression level of the genes. Such information is essential to further analyze a metagenomic data sets in-depth, for example, to define the origin of a sample or to improve environmental parameters for production reactors such as biogas reactors. Knowing whether an organism expresses a gene is a crucial element in deciding whether a more detailed analysis of that organism in the biotope where the sample was taken is necessary or not.

## Availability and future directions

MUFFIN is an ongoing workflow project that gets further improved and adjusted. The modular workflow setup of MUFFIN using Nextflow allows for fast adjustments as soon as future developments in hybrid metagenomics arise, including the pre-configuration for other workload managers. MUFFIN can directly benefit from the addition of new bioinformatics software such as for differential expression analysis and short-read assembly that can be easily plugged into the modular system of the workflow. Another improvement is the creation of an advanced user and wizard user configuration file, allowing experienced users to tweak all the different parameters of all the different software as desired.

MUFFIN will further benefit from different improvements, in particular by graphically comparing the generated MAGs via a phylogenetic tree. Furthermore, a convenient approach to include negative controls is under development to allow the reliable analysis of super-low abundant organisms in metagenomic samples.

MUFFIN is publicly available at https://github.com/RVanDamme/MUFFIN under the GNU general public license v3.0. Detailed information about the program versions used and additional information can be found in the GitHub repository. All tools used by MUFFIN are listed in the supplementary table S1. The Docker images used in MUFFIN are prebuilt and publicly available at https://hub.docker.com/u/nanozoo, and the GTDB formatted for sourmash(v2.0.0a10)^38^ usage is publicly available at https://osf.io/wxf9z/ and was created by C. Titus Brown (associate professor at UC DAVIS, http://ivory.idyll.org/blog/2019-sourmash-lca-db-gtdb.html).

## Acknowledgment

We want to thank Hadrien Gourlé and Moritz Buck for the valuable insights into metagenomic analysis and Annotation.

